# Latent gene network expression underlies partial re-evolution of a polyphenic trait in the worker caste of ants

**DOI:** 10.1101/2025.11.19.689334

**Authors:** Angelly Vasquez-Correa, Johanna Arnet, Travis Chen, Ehab Abouheif

**Affiliations:** Center for Evolutionary & Organismal Biology, Women’s Hospital, School of Medicine, Zhejiang University, Hangzhou, China; McGill University, Department of Biology, 1205 Avenue Docteur Penfield, Montreal, Quebec, H3A 1B1, Canada

**Keywords:** latent developmental potential, GRN, ocelli, ants, eye-antenna disc, evolutionary reversion, Formicinae, gene regulatory network, polyphenism, developmental plasticity

## Abstract

Polyphenisms––where alternative phenotypes develop from a single genome in response to environmental cues––are not only widespread in nature, but also occur at multiple levels of biological organization, from cells to individuals to societies. Polyphenism is thought to promote phenotypic diversification through the gain, loss, and re-evolution of alternative phenotypes. After the origin of a polyphenism, one of the alternative phenotypes often retains the developmental capacity to produce the ancestral trait, thereby permitting the other to evolve rapidly. Yet, little is known about the developmental processes underlying the re-evolution of polyphenic traits and how they may produce phenotypic diversification. Here we address this question by focusing on the caste polyphenism in ant societies, which produces a winged queen caste and a wingless worker caste in a single colony in response to environmental cues. We show, in a hyperdiverse group of ants, that a caste-specific trait called the ocelli (3 simple eyes on the dorsal head) is always present across queen castes but was lost and partially re-evolved multiple times, giving rise to novel patterns (1 ocelli) in the worker castes. Surprisingly, we discovered that a hidden (latent) expression of the ocelli gene regulatory network in worker castes that lost ocelli underlies the partial re-evolution of ocelli in this group. We therefore propose that latent developmental potentials may generally persist across polyphenic systems, including ant castes, and may facilitate the partial re-evolution of novel phenotypic patterns.

## Introduction

Polyphenism is a form of developmental plasticity where alternative phenotypes develop from a single genome in response to environmental cues (Nijhout, 2003).It is phylogenetically widespread feature of plants and animals that has evolved at different levels of biological organization (Hanna et al., 2024; West-Eberhard, 1989, 2003). For example, at the population level, the mouth form polyphenism in nematode worms produces alternative big tooth (omnivorous) or small tooth (bactivorous) mouth forms that develop in response to pheromones, crowding, salt concentration, temperature, and culturing substrate (Bento et al., 2010; Bose et al., 2012; Ragsdale et al., 2013; Werner et al., 2018). At the colony-level, caste polyphenism in eusocial insects produces morphologically differentiated queen and worker castes that develop in response to temperature and nutrition (Chandra et al., 2018; Evans & Wheeler, 2001; Korb, 2025; A. Rajakumar et al., 2024). And finally, at the cellular-level, polyphenism occurs within a multicellular individual, where a single genome gives rise to differentiated cell-types during development, such as between germline and somatic cells, in response to internal cues like morphogen gradients (Brunet & King, 2017; Davison & Michod, 2021; Devlin et al., 2023). Polyphenism has been proposed to promote, at the macroevolutionary scale, phenotypic diversification through the gain, loss, and re-evolution of alternative phenotypes (West-Eberhard, 2003). This is based on the idea that, once a polyphenic trait originates, one of the alternative morphs retains the capacity to produce the trait in the genome while the other is freer to evolve. This hypothesis has received support from a comparative study of mouth form polyphenism across 90 species of nematode worms showing a phylogenetic association between the gain and loss of alternative mouth form phenotypes and the phenotypic diversification of mouth parts (Susoy et al., 2015). Another supporting example is fat synthesis in parasitic wasps, which revealed an association between developmental plasticity and the loss and subsequent re-evolution of fat synthesis in one species (Peters et al., 2017; Visser et al., 2010, 2021). Yet, the underlying developmental and genetic processes facilitating the gain, loss and re-evolution of polyphenic traits remain poorly understood. (Forni et al., 2026; Sommer, 2020; West-Eberhard, 2003).

Here we address this question by focusing on caste polyphenism in the eusocial colonies of ants, which consists of a morphological division of labor between a winged reproductive queen caste and wingless non-reproductive worker caste in almost all 16,962 valid described ant species (AntWeb, 2026; Ward, 2014). The differential expression of polyphenic traits, such as wings, that develop in queens but not workers, are called “caste-specific” traits (Miura, 2005). It has been shown that polyphenic traits, including caste polyphenism in ants, are produced during development by the differential expression of highly conserved gene regulatory networks (GRN) in response to environmental cues (Abouheif & Wray, 2002; Béhague et al., 2018; Casasa et al., 2021; Davidson et al., 2023; Lenuzzi et al., 2023; A. Rajakumar et al., 2024; Vizueta et al., 2025). However, how the expression of these GRNs influences the evolution of caste-specific traits in ants remains unknown.

Here we focus on the ocelli, which are 3 small single-lens eyes on the dorsal head of most flying insects. Ocelli complement the function of the compound eyes by mediating orientation using polarised light and in the synchronization of daily activity (Buschbeck & Bok, 2023; Krapp, 2009). We investigate the evolution of ocelli in a hyperdiverse subfamily of ants (Formicinae), where they are universally present in the winged reproductive caste (queens and males) as 3 large ocellus that aid in mating flights and dispersal (Moser et al., 2004; Narendra et al., 2016). In contrast, ocelli in the wingless worker caste are evolutionarily labile, showing dramatic variation across species in the presence / absence or number of ocelli in the worker caste (Johnson & Rutowski, 2022; Narendra et al., 2016; Narendra & Ribi, 2017; Schwarz et al., 2011) (Figure 1A). In some species, adult workers completely lack ocelli, such as in *Camponotus floridanus*, while in others they are present and vary in number––there are species whose adult workers have all 3 ocelli or just 1 single ocellus, and these can be present in all or in only in a subset of workers (Figure 1A). For example, workers in *Cataglyphis bicolor* have all three ocelli, which function in light sensing and navigation, acting as a celestial compass that provides crucial directional information (Fent & Wehner, 1985) (Figure 1A). In contrast, all workers of *Polyrachis bihamata* have just a single medial ocellus (Hung, 1967) (Figure 1A), and in workers of *Dinomyrmex gigas*, a single medial ocellus evolved only in a subset of individuals in the worker caste called ‘soldiers’ (or major workers) with large heads, but are absent in other individuals called ‘minor workers’ with small heads (AntWeb, 2026) (Figure 1A). How this dramatic variation in ocelli in the worker caste of formicine ants has evolved remains poorly understood.

**Figure 1.**
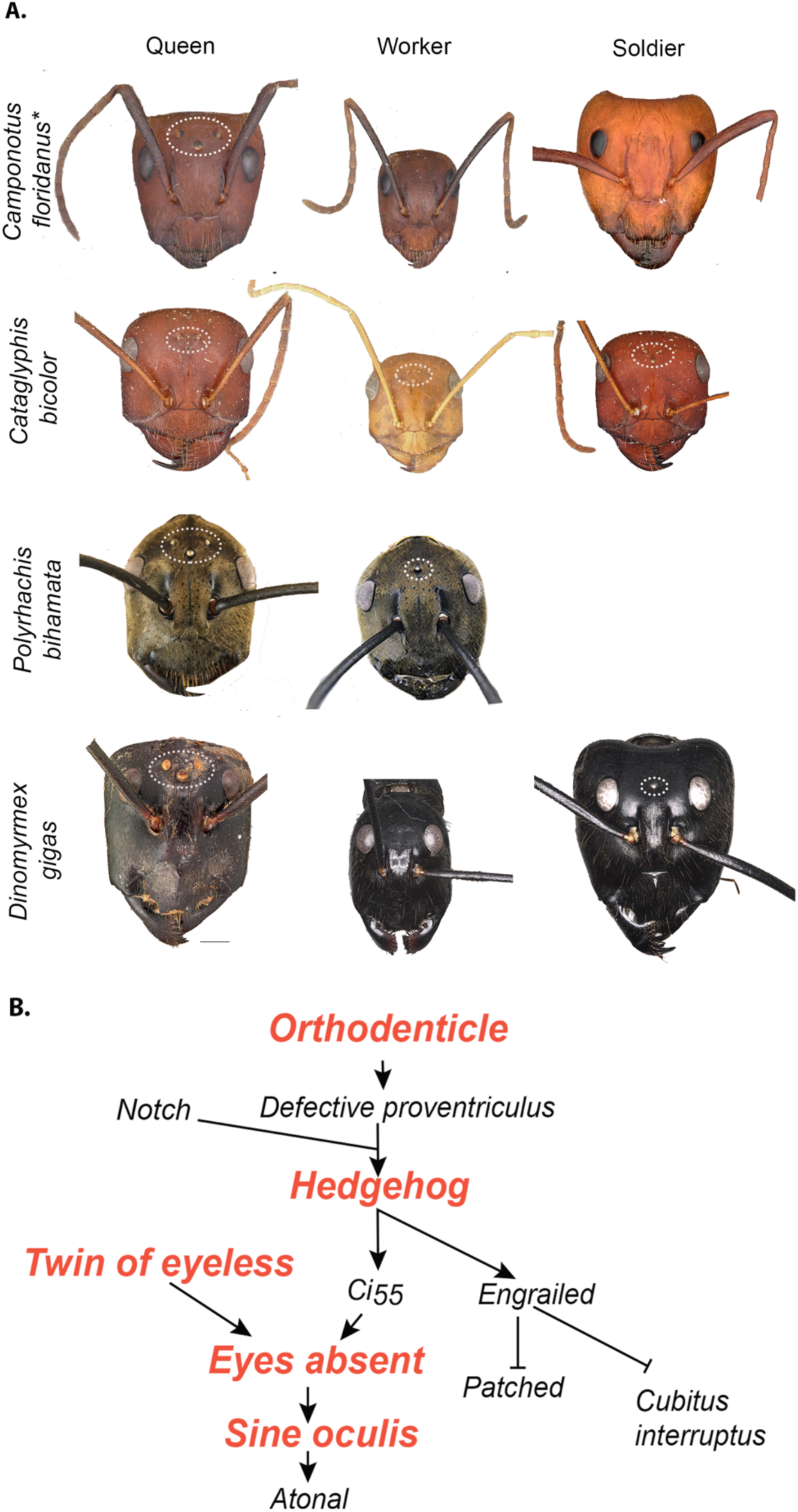
The presence and absence of ocelli in queens and workers across ants. (A) Ocelli develop in the winged reproductive castes across species of formicinae ants exemplified by *Camponotus floridanus*, *Cataglyphis bicolor, Polyrachis bihamata* and *Dinomyrmex gigas* (white dashed circles). Ocelli in the worker caste of formicine ants (white dashed circles) are evolutionarily labile, showing no ocelli (*Camponotus floridanus*), all 3 ocelli in all individuals of the worker caste (*Cataglyphis bicolor*) or only 1 ocellus in only the soldiers (*Polyrachis bihamata* and *Dinomyrmex gigas*). (B) Schematic representation of the ocelli GRN in *Drosophila melanogaster* adapted from Jean-Guillaume & Kumar (2022)(Jean-Guillaume & Kumar, 2022). The genes investigated in this study are highlighted in orange. Arrowheads indicate activation, and bars indicate repression. Queens are scaled to 1mm, and workers and soldiers are scaled according to the queen of each species. Asterisks indicate that *C. floridanus* was used for gene expression studies. Photos from Antweb (AntWeb, 2026)

This evolutionary lability of ocelli across the worker castes of formicine ants also provides an opportunity to understand how the GRN underlying development of ocelli influenced the evolution of this caste-specific trait. Ants are holometabolous insects, in which adult body parts develop from imaginal discs, semi-independent clusters of cells in the larvae (Held, 2002; Koch & Abouheif, 2019). In the fruit fly *Drosophila melanogaster*, ocelli develop from the eye-antenna imaginal disc located at the ventral region of the head capsule. The eye-antenna imaginal disc also gives rise to the head capsule, eye, antenna, and maxillary palps (Held, 2002). *D. melanogaster* is the only insect where the GRN underlying ocelli development has been well characterized at the third larval stage (Blanco et al., 2009; Domínguez-Cejudo & Casares, 2015; Sabat et al., 2017) (Figure 1B). The gene *orthodenticle* (*otd*) (formerly known as *ocelli-less*) is a selector gene that is necessary for specifying the ocellar region in the developing head capsule. *otd* expression in the ocellar domain, together with other genes like *hedgehog* (*hh*), initiates the development of the ocellar region and the three ocelli (two lateral ocellus and one medial ocellus) in the eye-antennal disc. The activation of these genes regulates the expression of downstream genes, such as *otd* regulating the expression of *defective proventriculus* (*dve)* (Blanco et al., 2009; Jean-Guillaume & Kumar, 2022; Yorimitsu et al., 2011), whereas *hedgehog* (*hh*) activates a portion of the retinal determination network, such as *eyes absent* (*eya*), *twin of eyeless* (*toy*), *sine oculis* (*so),* and *atonal* (*ato*). These retinal determination genes have been shown to be regulated by independent regulatory enhancers from the compound eye (Blanco et al., 2009, 2010; Jean-Guillaume & Kumar, 2022) (Figure 1B). Because the eye-antennae disc and the ocelli GRN have only been well characterized in *D. melanogaster*, it remains unknown whether they are conserved in ants.

To understand how the GRN underlying ocelli development may have influenced the evolution of this caste-specific trait, we first inferred the evolutionary history of ocelli in adult workers across the Formicinae using ancestral state reconstruction. We then characterized the eye-antennae disc in ants using three genes, *eyeless* (*ey*), *distal-less* (*dll*), and *otd-1*, which are known to mark the eyes (*ey*), the antenna (*dll*), and the head capsule and ocelli (*otd-1*). This characterization allowed us to investigate the expression of five key genes in the ocelli GRN, *otd-1, hh, toy, eya,* and *so*, during development of the winged and wingless castes across two formicine species.

## Results

### Partial reversion to a single ocellus occurs 3 times independently within the tribe Camponotini (Formicinae)

We first performed an ancestral state reconstruction to infer the evolutionary history of worker ocelli across the subfamily Formicinae. Our ancestral state reconstruction inferred a single re-gain of worker ocelli at the base or early within the Formicinae (Figure 2). Subsequent to the gain of ocelli early in the evolution of the Formicinae, our analysis inferred a single, well-supported, loss of worker ocelli at the base of the tribe Camponotini (Figure 2). Following this single loss, we inferred three (well supported) independent and partial re-evolution to a single medial ocellus in three different genera: *Camponotus gibbinotus*, *Polyrachis bihamata*, and *Dinomyrmex gigas* (Figure 2). In *P. bihamata*, the single ocellus occurs in all workers in the colony, whereas in *C. gibbinotus* and *D. gigas* the single ocellus occurs only in the large-headed soldiers (Figure 1). We therefore investigated the developmental role of the ocelli GRN underlying these independent partial reversions to a single medial ocellus in this tribe of ants.

**Figure 2.**
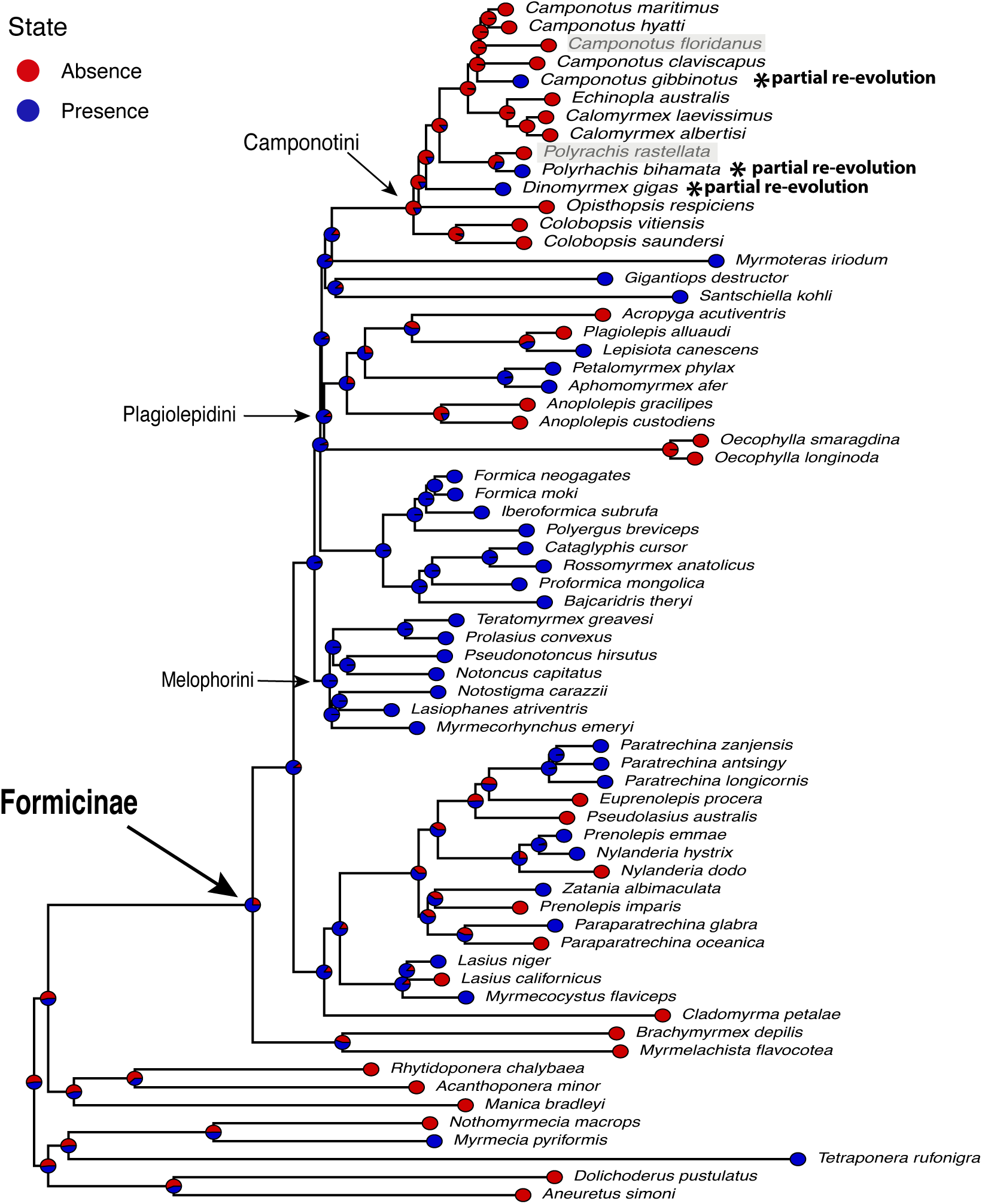
Ancestral state reconstruction of ocelli reveals three well-supported reversions of ocelli in the worker caste in the tribe Camponotini. Maximum clade credibility tree of formicine ants from Blaimer et al.(Blaimer et al., 2015). Ancestral state reconstruction for the presence (blue-colored circles) and absence (red-colored circles) of ocelli based on stochastic character mapping. Each pie chart for the nodes represents the posterior probabilities, scaled by the weight of evidence for each model. Species used to analyze ocelli GRN expression are highlighted in gray, and the species that re-evolved one ocellus is indicated as partial re-evolution. Three tribes within the Formicinae are marked (Melophorini, Plagiolepidini, Lasiini, Myrmelachistini and Camponotini) by arrows.

### A fate map characterizing the development of the head capsule, antennae, eyes, and ocelli within the eye-antenna disc in ants

To understand whether the GRN underlying the development of ocelli influenced the reversions of this caste-specific trait in workers, we first had to characterize the development of the eye-antenna disc in ants. In *D. melanogaster*, the eye, antennae, maxillary palps, ocelli, and head capsule develop from the eye-antenna disc, which is segregated into regions marked by the expression of highly conserved developmental genes (Haynie & Bryant, 1986; Held, 2002). In the Florida carpenter ant *C. floridanus*, we found that, similar to *D. melanogaster*, expression of *otd-1* marks the precursor regions of the head capsule and ocelli, *ey* marks the precursor regions of the eyes, and *dll* marks the precursor regions of the antennae. During the first larval instar, expression of *otd-1* emerges primarily in the head capsule in middle part of the disc between the antenna and compound eye (Figure 3B.). In contrast, *dll* and *eya* expression delineate the precursor regions of the antenna and eye (Figure 3C, D). During the second and third larval instar, *otd-1* is expressed in the developing head capsule and ocelli in the medial region of the disc (Figure 3 G, L), while *dll* is confined to the antenna and *ey* to the compound eye region (Figure 3H, I, M, N). Finally, during the fourth (final) larval instar, a developmental threshold mediated by juvenile hormone acts as switch point to determine whether larvae will develop either into a minor worker or soldier (MacMillan et al., 2025). Once larvae have been determined, expression patterns of *otd-1*, *dll,* and *ey* in worker-destined larvae (Figure 3 P to T) or soldier-destined larvae (Figure 3 U to Y) remain expressed in the same regions as in the second and third instars (Figure 3Q to T, V to Y). Together, our characterization shows that the eye-antennae disc and the regional identities within it, including the precursor region of the head capsule, ocelli, eyes, and antennae, are conserved in ants relative to *Drosophila*.

**Figure 3.**
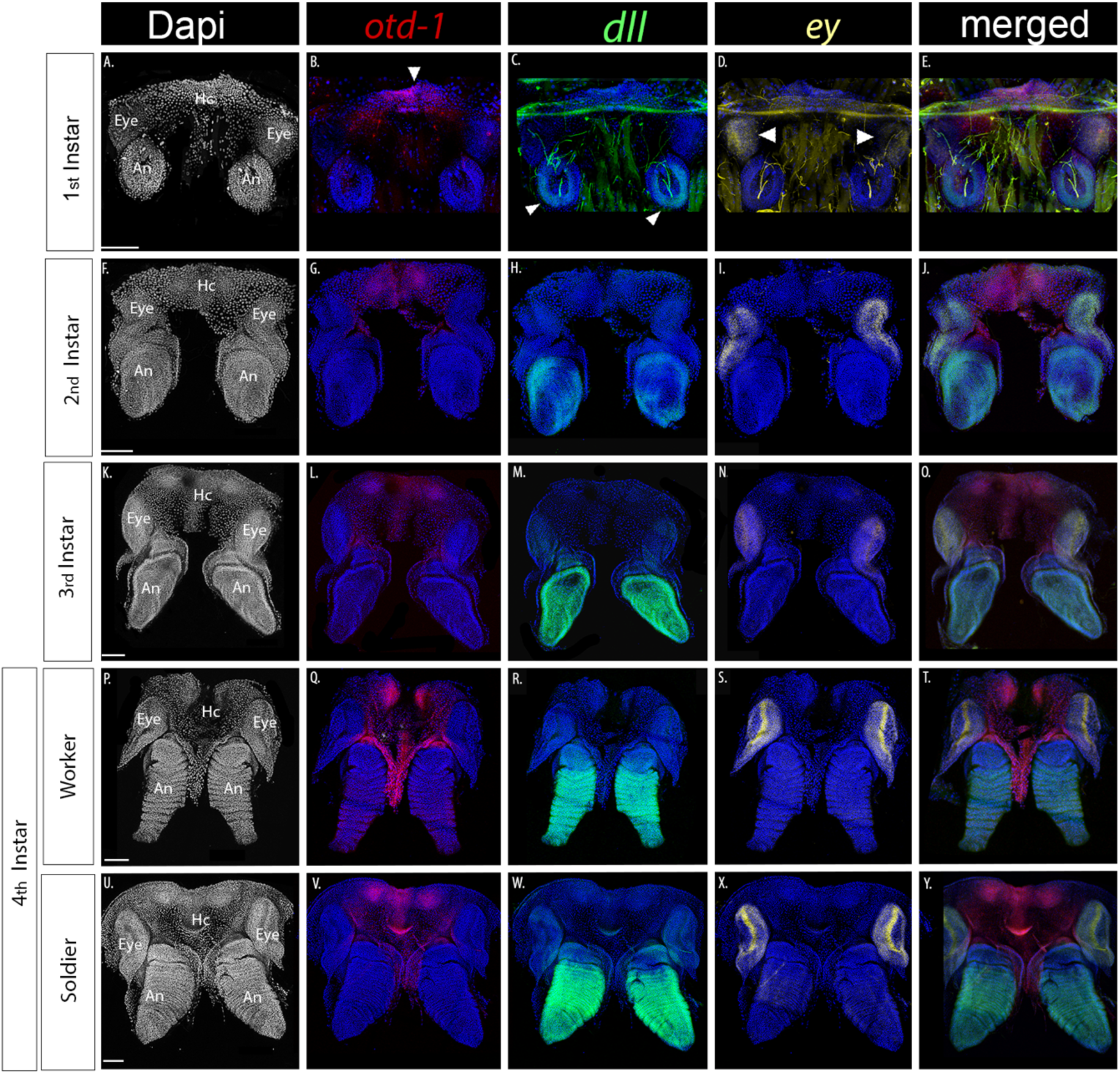
Characterizing development of the eye-antenna imaginal disc in worker castes of *C. floridanus* using *orthodenticle-1 (otd-1), distal-less (dll)* and *eyeless (ey)* gene expression to mark the developing head capsule and ocelli, antenna, and eyes. Fluorescent images in panels A, F, K, P, U represent the development of the entire eye-antenna imaginal disc marked with the nuclear stain DAPI across all four larval stages, where the head capsule region is labelled as ‘Hc’, the antennal region is labelled as ‘An,’ and the eye region is labelled as ‘Eye.’ Panels B, G, L, Q, V represent the development of the head capsule (Hc) marked by the genes *orthodenticle-1 (otd-1* in magenta); Panels (C,H,M,R,W) represent the antennal region marked by *distal-less* (*dll*) in green color; and Panels (D,I,N,S,X) represent the eyes (Eye) is *eyeless (ey*) (yellow); Panels. (A-E) First instar. Note: the green or yellow staining outside of the structures highlighted by the white arrows in panels C, D and E is background noise. (F-J) second instar, (K-O) third instar, (P-T) fourth instar worker-destined larvae and, (U-Y). Fourth instar soldier-destined larvae. All images are to scale.

### Expression of the ocelli GRN is conserved in winged reproductive castes but is latent in species whose adult workers completely lack ocelli

We next asked whether the ocelli GRN is conserved in the winged reproductive caste (males) relative to *Drosophila* and whether it is expressed in workers that entirely lack ocelli as adults. We address these questions using two species *C. floridanus* and *Polyrachis rastellata*. We chose these two species because *C. floridanus* is closely related to *C. gibbinotus* and *P. rastellata* is closely related to *P. bihamata*, which are two of the species that our ancestral state reconstruction inferred independent partial reversions to single medial ocellus in the worker caste (Figure 2). In the winged male caste of *C. floridanus*, we found that *otd-1, eya* and *so* are expressed where the 3 ocelli will develop, while *toy* and *hh* are expressed in the inter-ocellar region (the tissue that separates the three ocellus) (see white arrowheads in Figure 4A to B ’ and 5A to 5C’). Because these genes are similarly expressed within the ocellar region within the eye-antennal disc of *D. melanogaster*, we infer that the ocelli GRN is conserved in the winged reproductive castes in ants.

**Figure 4.**
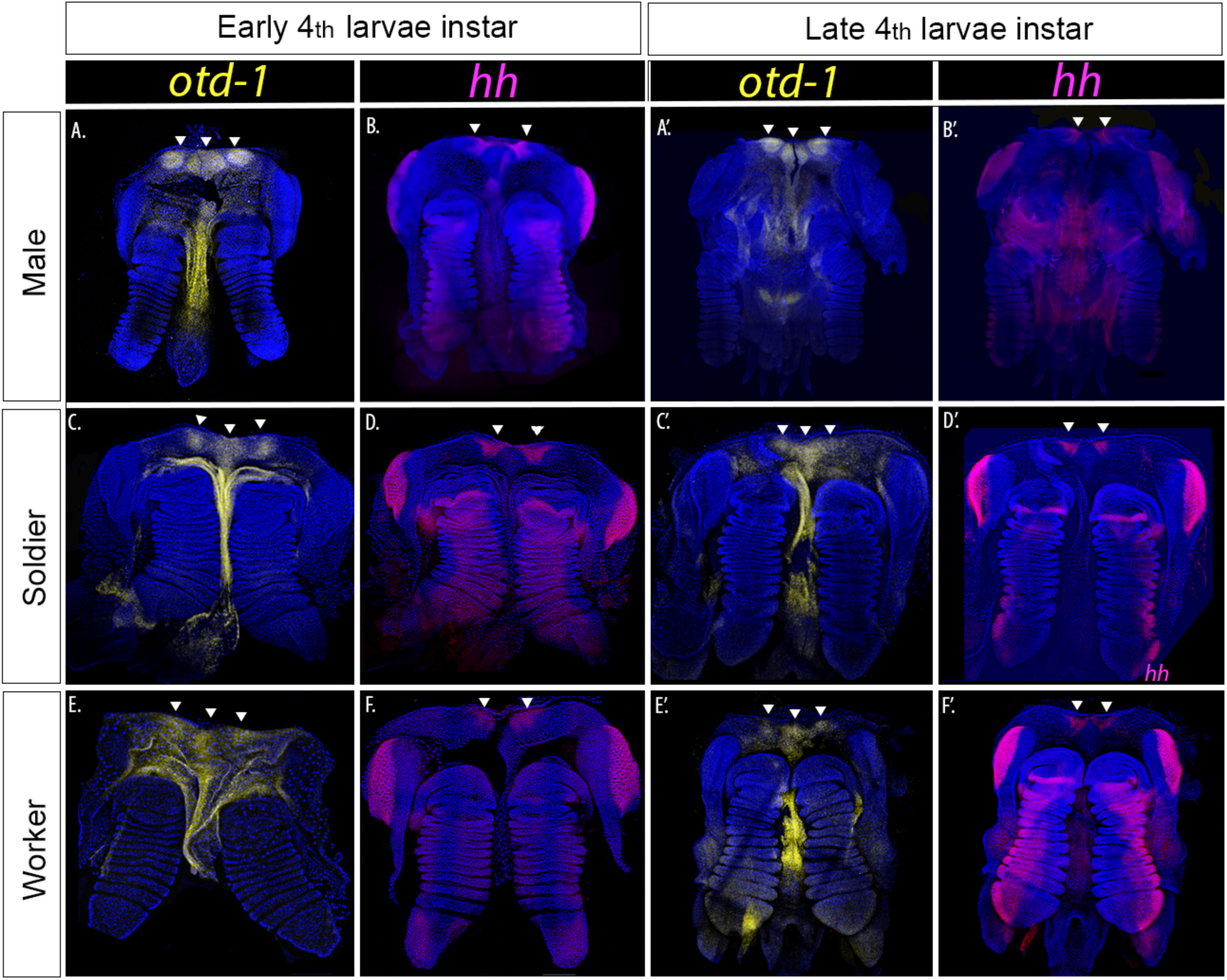
Latent expression of *otd-1* and *hh* genes in the ocelli GRN in workers and soldiers of *C. floridanus* during the 4^th^ larval instar. Expression of *orthodenticle-1(otd-1)* is yellow, and *hedgehog (hh)* is magenta. Early 4^th^ instar; (A, B) males (C, D) soldiers and (E, F) workers. Late 4^th^ instar: (A,’ B’) males (C’, D’) soldiers and (E’, F’) workers.

Surprisingly, we discovered that the ocelli GRN remains latently expressed in minor worker- and soldier-destined larvae of *C. floridanus*, which completely lack ocelli as adults. In soldier-destined larvae, all five genes remain expressed in the ocellar region within the eye-antennal disc at the beginning of the last larval instar (Figure 4C, D and Figure 5D to F). By the end of this instar, *otd-1* and *hh* remain expressed (Figure 4C’, D’), but *toy*, *eye*, and *so* are either down-regulated or absent relative to their expression in the compound eye region (Figure 5D’ to F’). In minor worker-destined larvae, 3 of the 5 genes (*otd-1*, *hh, eya*) remain expressed in the ocellar region during the early part of the last larval instar (Figure 4E, F and Figure 5 H), whereas *toy* and *so* are expressed in the compound eye region but absent (interrupted) in the ocellar region (Figure 5G, I). Furthermore, in *P. rastellata*, whose worker caste is composed of similarly sized individuals with no subcastes, we found that *otd-1*, *eya*, and *so* remain expressed in the ocellar region within the eye-antennal disc during the early part of the last larval instar (Figure 6). Together, our results show that despite the absence of ocelli in adult workers for millions of years, the expression of the ocelli GRN remains latent during larval development.

**Figure 5.**
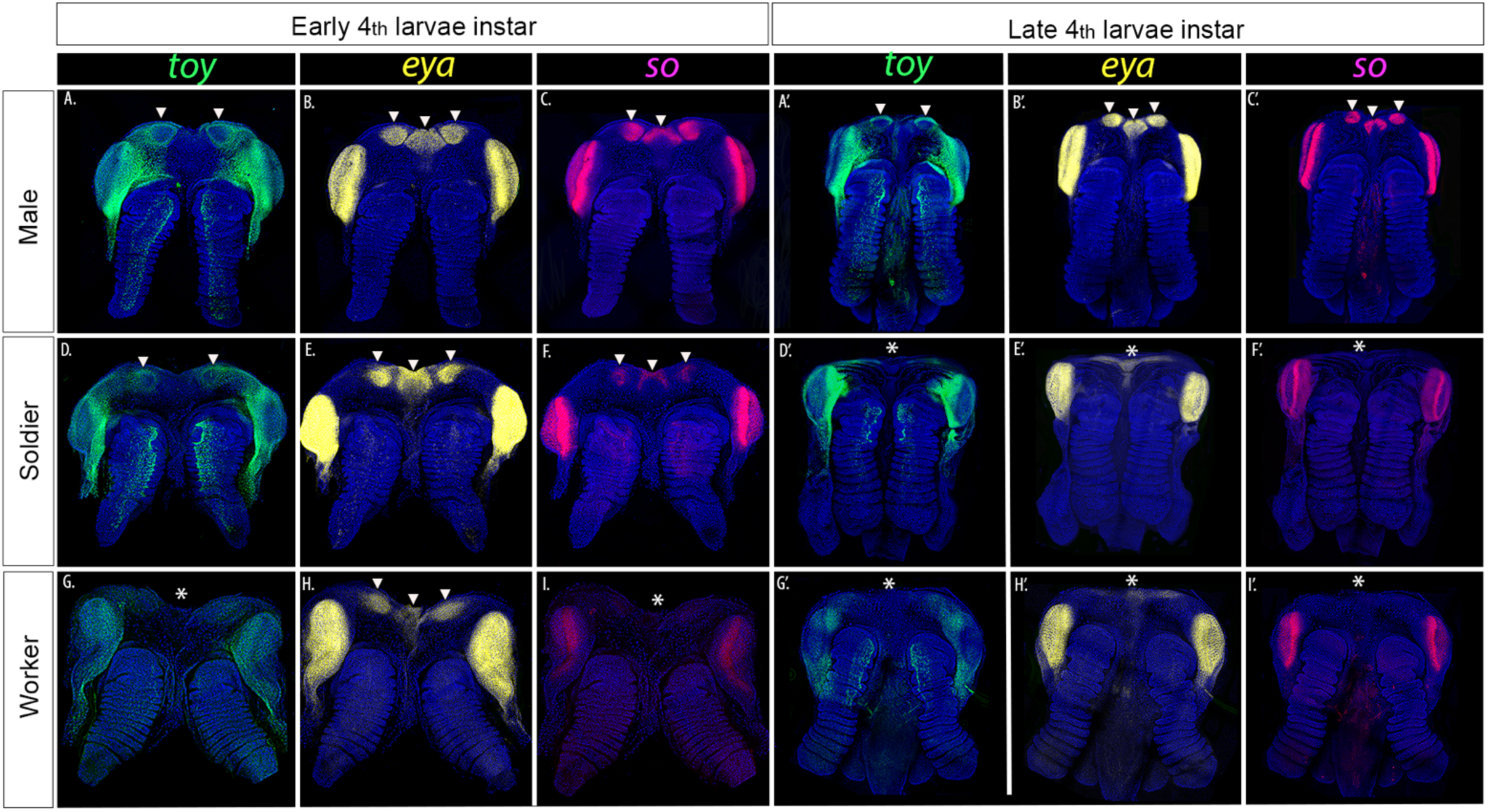
Latent Expression of *toy*, *eya*, and *so* within the ocelli GRN in the developing worker and soldiers of *C. floridanus*. Expression of *toy* (green), *eya* (yellow), and *so* (magenta). Early 4^th^ instar;(A to C) males, (D to F) soldiers and (G to I) workers. Late 4^th^ larvae stage;(A’ to C’) males, (D’ to F’), soldiers and (G’ to I’).

**Figure 6.**
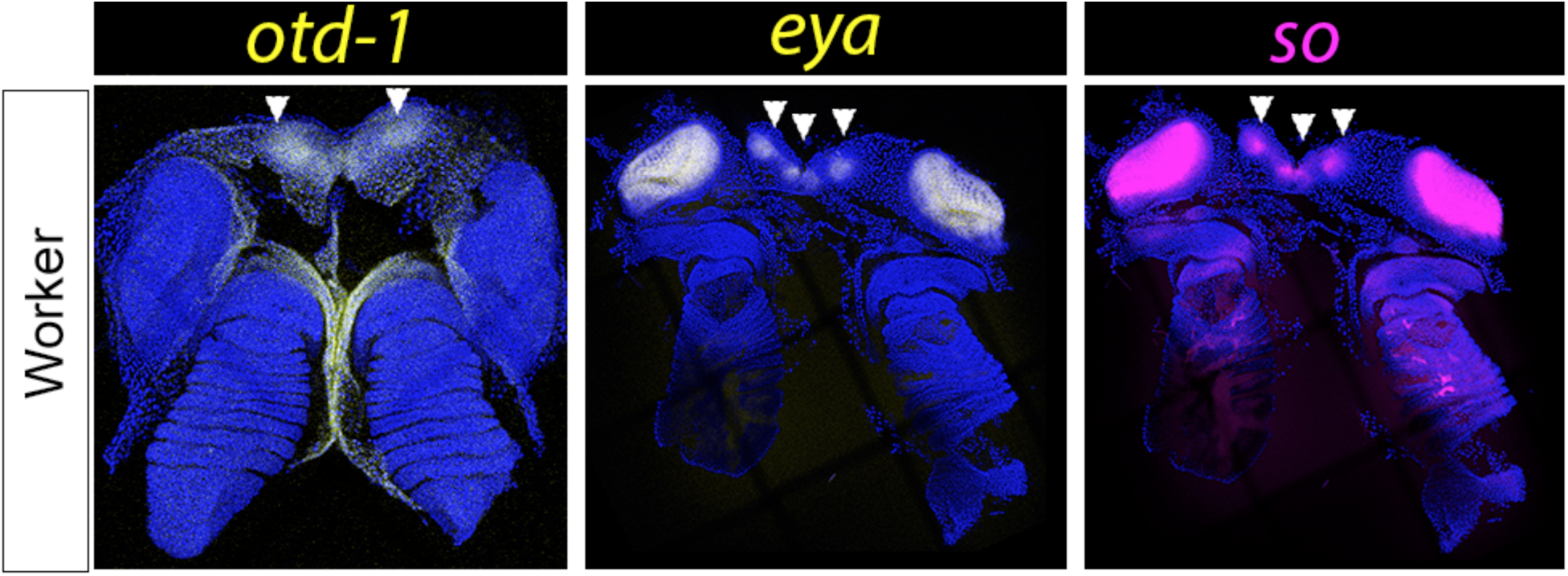
Latent expression of the ocelli GRN in the eye-antennal disc in worker larvae of *Polyrachis rastellata* at 4^th^ larvae stage. Expression of selected genes *otd-1* (yellow), *eya* (yellow), and *so* (magenta) at early 4th instar larvae.

Finally, we performed Scanning Electron Microscopy (SEM) in *C. floridanus* male, soldier, and minor worker pupae. In minor worker and soldier pupae, we discovered the existence of rudimentary ocelli that appear at the beginning of pupal development, continue to be elaborated, and then are eliminated before they molt into adult workers (Figure 7). These ocelli rudiments in are highly reduced relative to the fully functional ocelli found in male pupae. Finally, the pattern and timing of development of ocelli rudiments in the minor worker and soldier pupae coincides with the spatial expression and timing of interruption of the latent expression of the ocelli GRN. In soldier-destined larvae, ocelli GRN expression is interrupted later in development than in minor worker-destined larvae, and consequently, the ocelli rudiments in soldier pupae continue to develop longer and are more elaborated relative to those in minor worker pupae (Figure 7B to D and F to H). Therefore, expression of the latent ocelli GRN in the eye-antennae disc results in the development of ocelli rudiments in worker pupae and are then eliminated in adult workers.

**Figure 7.**
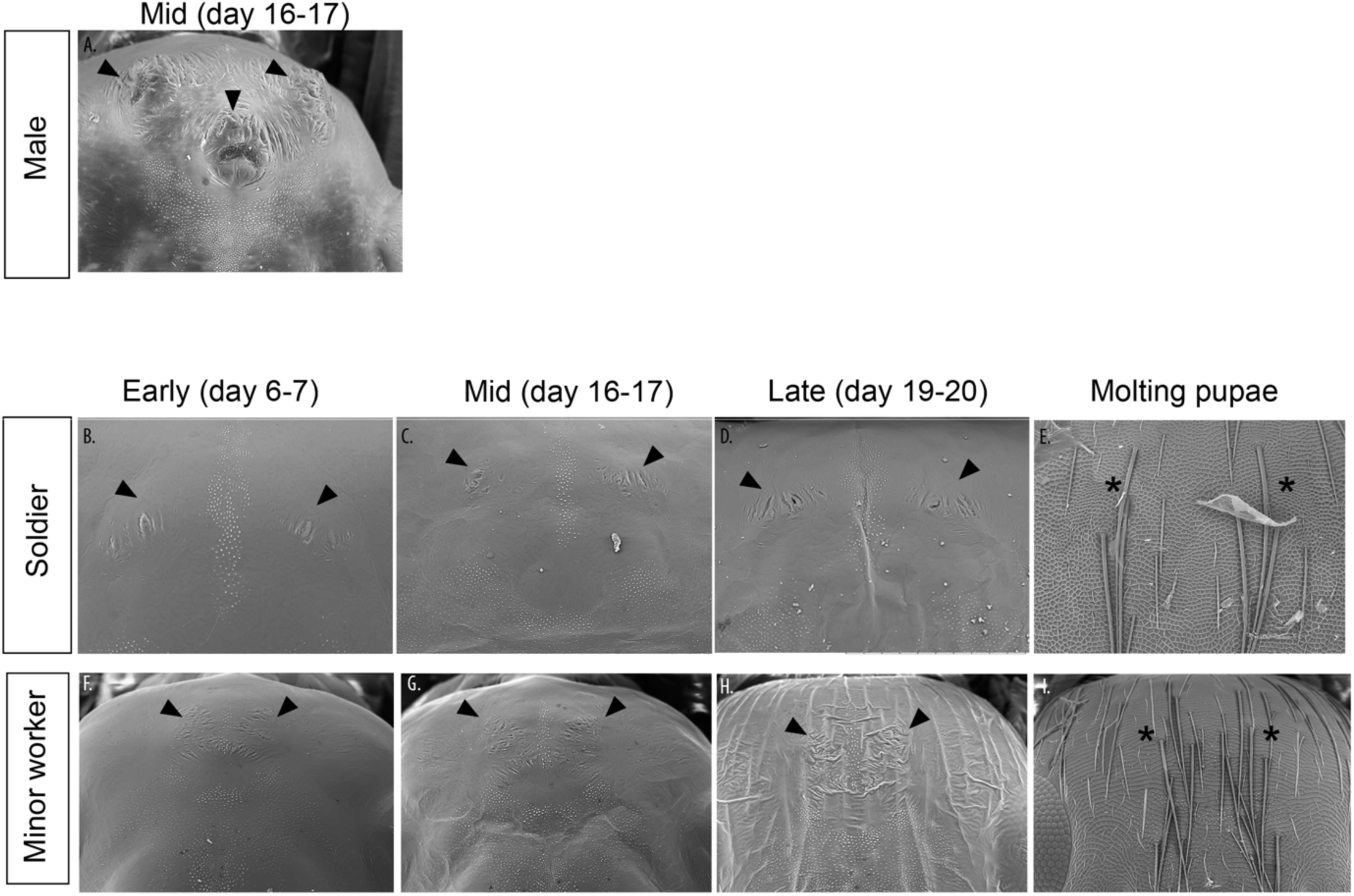
Development of rudimentary ocelli in worker and soldier pupae of *C. floridanus*. (A) SEM showing ocelli development in males at mid-stages of pupal development. SEM showing development of rudimentary ocelli on (B to E) soldiers and (F to I) minor workers during early (day 6-7), mid (day 16-17), and late (19-20) pupal development. These ocelli rudiments disappear prior to adult stage (E and I).

## Discussion

Our developmental and evolutionary data provide evidence that the latent expression of genes in workers lacking ocelli as adults is part of a latently expressed ocelli GRN, which likely facilitated at least 3 independent evolutionary reversions of this trait in the worker caste of species within the Camponotini clade. The latent expression patterns of genes in the ocelli region of developing workers lacking ocelli as adults are the same as in males that will develop fully functional ocelli but are only interrupted late in larval development. Furthermore, the timing and pattern of these late interruptions coincide with the degree of development of rudimentary ocelli in minor worker and soldier pupae before they disappear in adults (Figure 7). This indicates that, although the expression of these genes is latent, they still retain the capacity to produce rudimentary ocelli in the pupal stage before they disappear in adults. And finally, although *eya*, *toy*, and *so*, are part of both the ocelli and compound eye GRNs in *Drosophila*, the ocelli GRN has its own distinct identity and the genes within this GRN have distinct regulatory elements and are selectively regulated (Jean-Guillaume & Kumar, 2022; Zimmerman et al., 2000). In ants, our data shows that the selector gene for compound eye development in insects (*eyeless* / *Pax-6*) is expressed in the compound eyes and not ocelli, and the selector gene for ocelli (*otd-1*) is expressed in the ocelli and not compound eyes. We further show that the latent expression of *toy*, *eye*, and *so* are downregulated or absent in the ocellar region, but at the same time, are strongly expressed in the regions of the compound eyes. Therefore, the mutually exclusive expression of the selector genes *eyeless* / *Pax-6* in the compound eyes and *otd-1* in ocelli suggests that, like in *Drosophila*, the compound eye and ocelli GRNs have distinct identities, ultimately leading to differential expression of downstream genes and production of different cell types; compound eyes are produced from multiple imaging forming facets, while ocelli are produced from a single lense (Buschbeck & Bok, 2023; Jean-Guillaume & Kumar, 2022; Mishra et al., 2021). Altogether, our data show that the expression of these genes in workers lacking ocelli as adults are part of a latently expressed ocelli GRN.

Several hypotheses may explain how the ocelli GRN came to be latently expressed and maintained in developing workers that lack ocelli as adults. Perhaps the most simplistic hypothesis proposes that the presence of functional ocelli in adult queen and male castes maintains the ocelli GRN intact in the genome by keeping it under positive natural selection. This hypothesis assumes that this, as a side consequence, leads to expression of the ocelli GRN in the worker castes lacking ocelli. However, caste determination between queens and workers occurs through the action of a developmental threshold or switch, where a continuous environmental cue is translated into discrete phenotypic outcomes (Abouheif, 2021; MacMillan et al., 2025; Qiu et al., 2022; A. Rajakumar et al., 2024; Schultner et al., 2023). Once caste determination has occurred, the genome is expressed differentially during the developmental trajectories of queens and workers (Abouheif, 2021; Abouheif & Wray, 2002; Barkdull & Moreau, 2023; Béhague et al., 2018; Chandra et al., 2018; Khila & Abouheif, 2010; Qiu et al., 2022; Vizueta et al., 2025). These trajectories are decoupled, and consequently, can evolve largely independently (Abouheif, 2021; Powell et al., 2020; Vong et al., 2025). The dramatic variation in the number, size, and presence/absence of ocelli in worker castes across the Formicinae (see Figure 1) supports the largely independent evolution of ocelli in workers from those in queens, which always develop 3 ocellus. Furthermore, we observe similar patterns of variation in the wings and ovaries between queen and worker castes across ant species (Cronin et al., 2013; Khila & Abouheif, 2010; Monnin & Peeters, 2008; R. Rajakumar et al., 2012, 2018). Therefore, while the presence of ocelli in males and queens maintains the ocelli GRN in the genome and creates a potential for expression of this GRN in developing workers lacking ocelli, this cannot solely explain how this latent expression became actualized (released and fixed) and what maintained it over millions of years.

One hypothesis for how this ocelli GRN became latently expressed and has been maintained in workers lacking ocelli is pleiotropy, which potentially results from the multiple roles that genes within the ocelli GRN play within the same imaginal disc (the eye-antennal disc). For instance, *otd-1* and *hh* also determine the regional identity of the head capsule, while *hh*, *toy*, *eya* and *so*, also play key roles in compound eye development in specifying structures such as optic lobes, cone differentiation, and rhabdomeres development (Blanco et al., 2009; Domínguez-Cejudo & Casares, 2015; Jean-Guillaume & Kumar, 2022). Another example is the highly conserved developmental regulatory gene *sonic hedgehog* (*shh*), which plays a key role in limb development across animals. The vestigialization of hindlimbs in snakes and therefore the re-evolution of hindlimbs in extinct species across the phylogeny of the group, is thought to be maintained during development through pleiotropic enhancers that drive *shh* expression. This means that the same enhancer (ZRS) that drives *shh* expression in the external genital, also drives it (pleiotropically) in the developing limb region of snakes (Leal & Cohn, 2018). Future studies should attempt to explore whether expression of these conserved genes in multiple regions of the eye-antennal disc is driven by shared enhancers. Alternatively, we cannot rule out the hypothesis that ocelli GRN has been co-opted to play a novel, yet currently unknown, function during worker larval development. Recent discoveries on the evolution of the wing GRN in ants provides support for this hypothesis. The wings, another nearly universal caste-specific trait in ants, develop in the reproductive male and queen caste, but are halted in the worker caste in response to environmental cues (Abouheif & Wray, 2002). The wing GRN is also found to be latently expressed in the wingless worker caste of ants and was thought to be functionless (Abouheif & Wray, 2002). However, it was recently discovered that the latent expression of this wing GRN in wingless worker caste acquired a novel function to generate big-headed soldiers in the hyperdiverse ant genus *Pheidole* (R. Rajakumar et al., 2012, 2018).

Finally, our inference that this latent expression of the ocelli GRN in workers facilitated the partial reversion to a single medial ocellus is supported by: (1) the close phylogenetic relationship between species that lack ocelli in adult workers but retain a latent expression of the ocelli GRN (*C. floridanus* and *P. rastellata*), and those that underwent a partial phylogenetic reversion to a single medial ocellus (*C. gibbinotus*, *D. gigas* and *P. bihamata*); (2) the presence of a developmental capacity or potential of the latent ocelli GRN expression to produce rudimentary ocelli in the pupal stage of *C. floridanus* workers that completely lack ocelli as adults; (3) the ability to experimentally induce only 1, only 2, or all 3 ocelli in similarly sized adult workers that normally lack them by applying high doses of Juvenile Hormone (JH) to worker-destined larvae in the ant *Monomorium pharaonis* (Li et al., 2024); and finally (4) in nature, the rare induction of a single medial ocellus by mermithid parasites in soldiers of *Pheidole pallidula* that typically lack ocelli in natural colonies (Laciny et al., 2019; Passera, 1976). The natural or experimental induction of worker individuals with only a single medial ocellus in different ant species also supports the inference that the single medial ocellus can be developmentally dissociated from the other two lateral ocelli. This suggests that reversion can facilitate the appearance of novel patterns of ocelli development in the workers if selected for.

In *C. floridanus* and *P. rastellata*, there is a latent ocelli GRN expression for all 3 ocelli, providing a springboard to facilitate the partial phylogenetic reversion to a single medial ocellus in *C. gibbinotus*, *P. bihamata* and *D. gigas*. In the genus *Polyrhachis*, however, some species in the same subgenus as *P. bihamata*, such as *P. bellicosa*, have three ocelli. Because the phylogenetic relationships within this subgenus have yet to be resolved, the independent reversion of ocelli in the ancestor of this subgenus may have resulted either in a single ocellus as reflected in *P. bihamata* or in 3 ocelli as reflected in *P. bellicosa* and 2 ocelli were subsequently lost giving rise to the single medial ocellus in *P. bihamata* (Hung, 1967). These possibilities further reinforce the different evolutionary pathways by which this latent potential may facilitate novelty after reversion.

Future functional, genomic, and comparative analyses of the ocelli GRN between larval stages, individuals within the worker caste, and species will ultimately reveal the architecture of the ocelli GRN and whether its underlying enhancers and promoters are modular or pleiotropic. This, in combination with manipulations of insect hormones, such as JH and ecdysone, would also elucidate whether variation in the size, presence/absence, number of ocelli is regulated by continuous or switch-like developmental mechanisms. And finally, determination the organismal and ecological function of the ocelli will be important to understand the adaptive significance of the latent expression of the ocelli GRN at both the individual and colony-level.

More broadly, our findings suggest that the ancestral and latent GRN expression (also known as ancestral developmental potential) we observed may generally underlie polyphenic systems, including caste-specific traits in ants and other eusocial organisms. We therefore propose that ancestral developmental potentials facilitate the re-evolution of polyphenic traits (West-Eberhard, 2003), and when these potentials facilitate only the partial re-evolution of alternative phenotypes, novel phenotypic patterns appear. We hope our findings not only inspire future work testing these proposals in polyphenic organisms, but also in non-polyphenic ones, where the polyphenism occurs at the cellular level but not at the level of the whole organism. Here, the cellular polyphenism produces alternative cell-types from a single genome in response to internal cues, such as morphogen gradients within the organism. If alternative cell types retain homologs or serial homologs of specific trait (Jackman et al., 2025; V. J. Lynch, 2023), then this raises the possibility that ancestral and latent developmental potentials may generally facilitate re-evolution of alternative cell types in multicellular organisms.

## Data and code availability

All data reported in this paper are provided in Table S2 in the supplemental information. This paper does not report original code. Any additional information required to reanalyze the data reported in this paper is available from the lead contact upon request.

## Acknowledgments

We thank Hermogenes Fernandez-Marín, who inspired this work by sharing his knowledge about ocelli in soldiers across the *Atta* species. We thank Lloyd Davis, Marc Seid, Rajendhran Rajakumar, and Shelly Berger for help with collecting *Camponotus floridanus* colonies. We thank Mary Jane West-Eberhard, Friedrich Markus, Guilherme Gainett, Arjuna Rajakumar, and Rajendhran Rajakumar for discussions and/or comments on the manuscript, and Juan Carlos Penagos for input on phylogenetic analysis. We thank Erik Plante and undergraduates for help with feeding and maintaining lab colonies of *Camponotus floridanus*. Finally, we thank McGill University’s Integrated Quantitative Biology Initiative (IQBI) and Advanced Bioimaging Facility (ABIF) for imaging support. This work was supported by a Natural Sciences and Engineering Research Council of Canada (NSERC) Discovery Grant to E.A and by a Doctoral fellowship from NSERC BESS-CREATE program to A.V-C.

## Author Contribution

AV-C and EA conceived the project. AV-C and JA gathered ocelli data. AV-C and JA conducted ancestral state reconstruction. A V-C conducted HCR. TC performed SEM imaging. AV-C and EA wrote the manuscript.

## Methods

### Ant maintenance and collection

Colonies of *C. floridanus collected at Gainesville* (*Florida,* USA) and *P. rastellata* collected at Mae Tang (Chiang Mai, Thailand), were maintained in plastic boxes with glass test tubes filled with water-constrained cotton wool. They were fed mealworms and the Bhatkar–Whitcomb diet (Bhatkar & Whitcomb, 1970). Colonies were maintained at 25°C with 60% humidity in complete darkness.

### Larvae fixation and *in-situ* HCR

Gene sequences were obtained from NCBI GenBank database(Sayers et al., 2022) using genome BLAST against the assembled *C. floridanus* genome: *eyeless (ey*; XM_025414466), *distal-less (dll*; XM_025412727.1), *hedgehog, (hh*; XM_011262474.3), *eye absent (eya*; XM_025414466), *sine oculis (so;* XM_011252868.3) and *twin of eyeless (toy*; XM_011268499.3) genes. For *otd*, two paralogs of the gene were found in ants (XM_020028684.2 and XM_025415314.1), which is a result of a gene duplication event that has also been reported in wasps, bees, and beetles (J. A. Lynch et al., 2006; The Honeybee Genome Sequencing Consortium, 2006; Zattara et al., 2017). The two *otd* paralogs sequences were then aligned by multiple sequence alignment using all of the known *orthodenticle* related sequences in insects; *Drosophila melanogaster* (NM_001369965.1), *Apis mellifera*: *otd-1* (XM_026446161.1), *otd-2*(XM_006571236.3), *Nasonia vitripennis*: *otd-1* (XM_008212114.4), *otd-2* (XM_031926951.2) and *Tribolium castaneum otd-1* (XM_008192467.2), *otd-2* (XM_008192470.2) and *Acythosiphon pisum* (XP_008180802.1). To determine the *otd-1* paralog to *Drosophila melanogaster* (*otd-1)* a maximum likelihood gene tree was inferred using genetic distance model Hasegawa Kishino Yano (HKY) and 500 bootstrap replicates as incorporated in MEGA12 alpha (Kumar et al., 2024)(Figure S1). Probes corresponding to all genes were chosen for the hybridization chain reaction experiments using the fluorescence Hairpins (B1 546, B2 488, B3 647) synthesized by Molecular Instruments.

First, second, third and fourth larval instars of soldier, and minor worker-destined larvae and fourth instar of male destined larvae of *C. floridanus* and worker larvae of *P. rastellata* were collected and subsequently fixed in a PEM 4% formaldehyde solution for 2 hrs at room temperature. Fixed samples were then dehydrated progressively in methanol baths (25%, 50%, 75% methanol for 15 min each, and 100% overnight at 4°C) and stored in 100% methanol at −30°C until use. All gene expression analyses were conducted by In situ Hybridization Chain Reaction (HCR), following the protocol for HCR (v3.0 protocol) (Schwarzkopf et al., 2021). After the tissue was pre-hybridized in a prewarmed Probe Hybridization Buffer (Molecular Instruments) for 30 minutes at 37°C and incubated with HCR probes in a Probe Hybridization Buffer overnight at 37°C. Tissues were washed the next day in a prewarmed Probe Wash Buffer four times, 15 minutes each and washed in 5X SSCT (UltraPure 20XSSC Buffer, Invitrogen, diluted in water) three times for 5 minutes at room temperature. Tissues were pre-amplified in Amplification Buffer (Molecular Instruments) for 30 minutes at room temperature and incubated with snap-cooled HCR hairpins in Amplification Buffer overnight at room temperature. Tissues were then washed with 5X SSCT at room temperature twice for 5 minutes, for 30 minutes, and once for 5 minutes before being mounted on glycerol-DAPI 80%.

### Microscopy

Confocal imaging was used to describe gene expression using Leica SP8 confocal microscope. Fiji (Schindelin et al., 2012) was used for image processing. Scanning electron microscopy (SEM) was done on a Hitachi TM3030 Scanning Electron Microscope.

### Evolution of Ocelli in the Formicine Clade

The evolution of ocelli on workers across the subfamily Formicinae was inferred using ancestral reconstruction (ASE) for discrete traits incorporated in the R package Phytools 4.3.3(Revell, 2024). The ASE analysis was based on the UCE70 phylogeny for the cade Formicinae published by Blaimer *et.al.* (Blaimer et al., 2015). The species *Camponotus floridanus* and *Polyrachis bihamata* were added manually to the phylogeny. To determine the presence or absence of ocelli in workers, photographs of the studied species from the database AntWeb,Version 8.114 were used (AntWeb, 2026). The observations from the database were contrasted with published information from the literature (Table S2). Ocelli were classified as present in the worker caste if individuals exhibit any of the 3 ocellus (2 lateral and 1 medial ocellus). In the case of the presence of worker polymorphism, ocelli were classified as present if any one of the 3 ocelli was present within any of the worker subcastes. Whereas the absence was the complete lack of ocelli across workers and soldiers.

Four separate models of ocelli evolution were tested for each character in phytools: Equal rate “ER”, all transitions rate different “ARD”, and an irreversible model allowing only transitions between presence and absence, and another irreversible model allowing only transitions between absence and presence. We compared the fit of our models by computing Akaike information criterion (AIC) and Akaike weights and conducting pairwise likelihood ratio tests. The new function incorporated in phytools 4.3.3, *simmap,* was used to generate stochastic character maps under each of the four models tested (see Table S1). The stochastic mapping that resulted from the stochastic simulation represented the frequencies that are equal to the weight of evidence supported by each model (Revell, 2024)

## Supplementary Figures

**Supplementary Figure 1.**
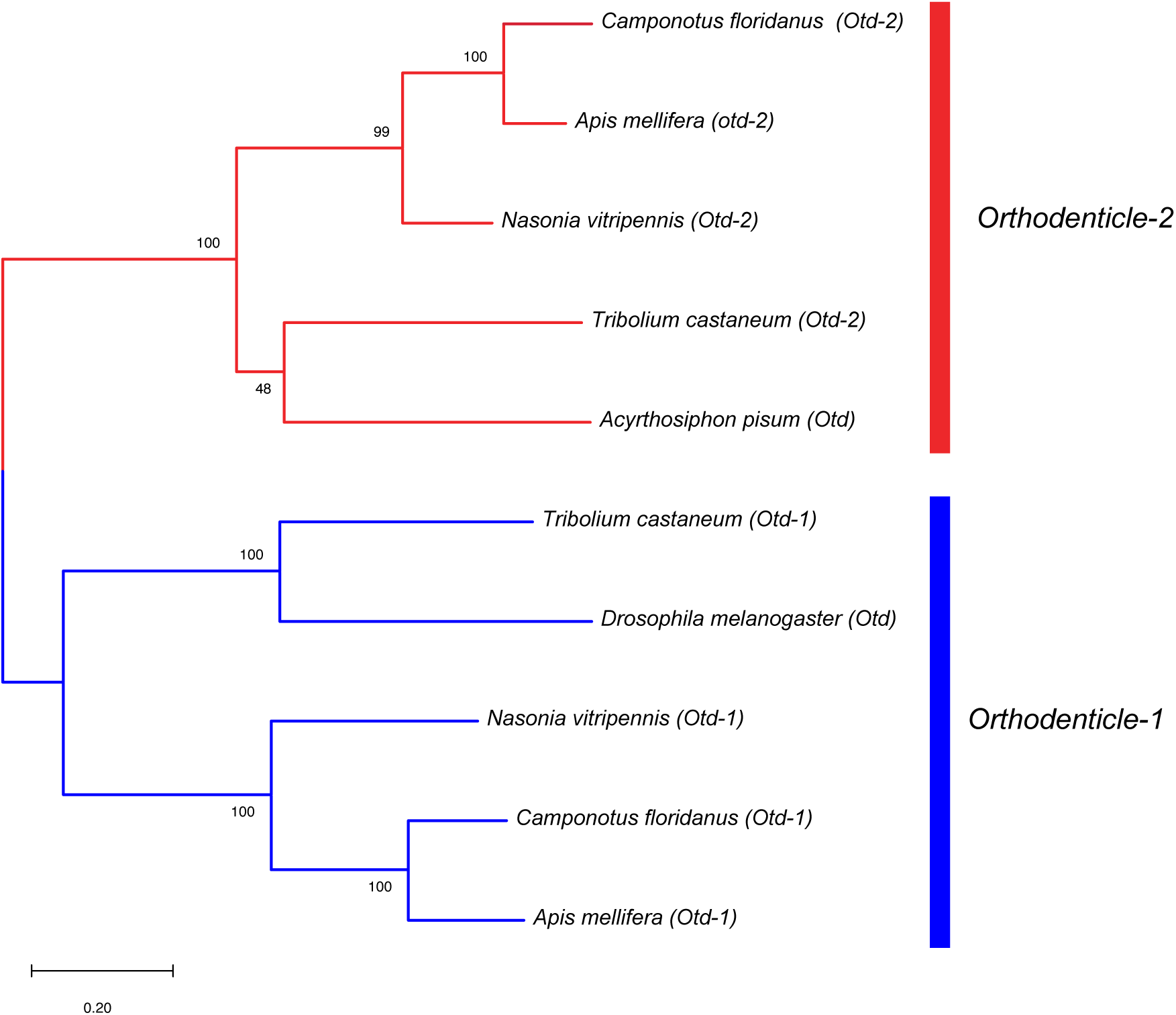
Simplified Gene tree based on maximum likelihood showing relationships between *orthodenticle (otd)* orthologs in *Drosophila melanogaster*, *Apis mellifera*, *Nasonia vitripennis*, *Tribolium castaneum, Acythosiphon pisum and Camponotus floridanus.* Branch values are bootstrap support (%). Colors: *otd-2* (red), *otd-1* (blue)

## Supplementary Tables

**Supplementary Table1.**
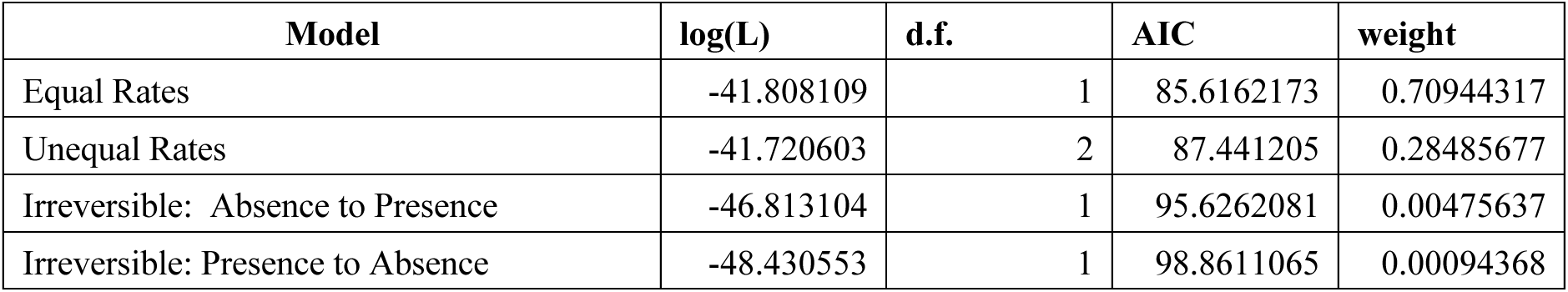
Model selection under maximum likelihood estimation implemented in phytools (Revell, 2024). The results are ordered by decreasing Akaike Weights (w).

**Supplementary Table 2.** Database with the references used in this study for presence and number of ocellus (1) and absence (0) of ocellus.

| Subfamily | Genus | Species | Ocelli Count | References |
| --- | --- | --- | --- | --- |
| outgroup | *Acanthoponera* | *minor* | 0 | AntWeb.org |
| Formicinae | *Acropyga* | *acutiventris* | 0 | AntWeb.org |
| outgroup | *Aneuretus* | *simoni* | 0 | AntWeb.org |
| Formicinae | *Anoplolepis* | *custodiens* | 0 | AntWeb.org |
| Formicinae | *Anoplolepis* | *gracilipes* | 0 | AntWeb.org |
| Formicinae | *Aphomomyrmex* | *afer* | 3 | AntWeb.org |
| Formicinae | *Bajcaridris* | *theryi* | 3 | Santschi, 1936 |
| outgroup | *Brachymyrmex* | *depilis* | 0 | AntWeb.org |
| Formicinae | *Calomyrmex* | *albertisi* | 0 | AntWeb.org |
| Formicinae | *Calomyrmex* | *laevissimus* | 0 | AntWeb.org |
| Formicinae | *Camponotus* | *floridanus* | 0 | AntWeb.com |
| Formicinae | *Camponotus* | *gibbinotus* | 1 | AntWeb.org |
| Formicinae | *Camponotus* | *hyatti* | 0 | MacKay & Mackay, 2002 |
| Formicinae | *Camponotus* | *maritimus* | 0 | Ward, 2005 |
| Formicinae | *Colobopsis* | *saundersi* | 0 | AntWeb.org |
| Formicinae | *Colobopsis* | *vitiensis* | 0 | Mann, 1920 |
| Formicinae | *Cataglyphis* | *cursor* | 3 | AntWeb.org |
| outgroup | *Cladomyrma* | *petalae* | 0 | Agosti, 1991 |
| outgroup | *Dolichoderus* | *pustulatus* | 0 | AntWeb.org |
| Formicinae | *Dinomyrmex* | *gigas* | 1 | AntWeb.org |
| Formicinae | *Echinopla* | *australis* | 0 | AntWeb.org |
| Formicinae | *Euprenolepis* | *procera* | 0 | Lapolla, 2009 |
| Formicinae | *Formica* | *moki* | 3 | Cole Jr, 1943 |
| Formicinae | *Formica* | *neogagates* | 3 | AntWeb.org |
| Formicinae | *Gigantiops* | *destructor* | 3 | Smith, 1858 |
| Formicinae | *Iberoformica* | *subrufa* | 3 | Antwiki- genus |
| Formicinae | *Lasiophanes* | *atriventris* | 3 | AntWeb.org |
| Formicinae | *Lasius* | *californicus* | 0 | AntWeb.org |
| Formicinae | *Lasius* | *niger* | 3 | AntWeb.org |
| Formicinae | *Lepisiota* | *canescens* | 3 | Sharaf et al., 2020 |
| outgroup | *Manica* | *bradleyi* | 0 | AntWeb.org |
| outgroup | *Myrmecia* | *pyriformis* | 3 | AntWeb.org |
| Formicinae | *Myrmecocystus* | *flaviceps* | 3 | AntWeb.org |
| Formicinae | *Myrmecorhynchus* | *emeryi* | 3 (*soldiers and media, absent in minors*) | Wheeler, 1917 |
| outgroup | *Myrmelachista* | *flavocotea* | 0 | AntWeb.org |
| Formicinae | *Myrmoteras* | *iriodum* | 3 | AntWeb.org |
| outgroup | *Nothomyrmecia* | *macrops* | 0 | AntWeb.org |
| Formicinae | *Notoncus* | *capitatus* | 3 | AntWeb.org |
| Formicinae | *Notostigma* | *carazzii* | 3 | AntWeb.org; Emery, 1920 |
| Formicinae | *Nylanderia* | *dodo* | 0 | LaPolla et al., 2011 |
| Formicinae | *Nylanderia* | *hystrix* | 3 | Kallal & LaPolla, 2012 |
| Formicinae | *Oecophylla* | *longinoda* | 0 | AntWeb.org |
| Formicinae | *Oecophylla* | *smaragdina* | 0 | Cole & Jones, 1948 |
| Formicinae | *Opisthopsis* | *respiciens* | 0 | AntWeb.org |
| Formicinae | *Paraparatrechina* | *glabra* | 3 | AntWeb.org |
| Formicinae | *Paraparatrechina* | *oceanica* | 0 | AntWeb.org |
| Formicinae | *Paratrechina* | *antsingy* | 3 | LaPolla & Fisher, 2014 |
| Formicinae | *Paratrechina* | *longicornis* | 3 | AntWeb.org |
| Formicinae | *Paratrechina* | *zanzensis* | 3 | LaPolla et al., 2013 |
| Formicinae | *Petalomyrmex* | *phylax* | 3 | Snelling, 1979 |
| Formicinae | *Plagiolepis* | *alluaudi* | 0 | AntWeb.org |
| Formicinae | *Polyergus* | *breviceps* | 3 | AntWeb.org; Smith, 1947 |
| Formicinae | *Polyrhachis* | *decumbens* | 0 | Kohout, 2006 |
| Formicinae | *Prenolepis* | *emmae* | 3 | AntWeb.org |
| Formicinae | *Prenolepis* | *imparis* | 0 | Williams & LaPolla, 2016 |
| Formicinae | *Proformica* | *mongolica* | 3 | AntWeb.org |
| Formicinae | *Prolasius* | *convexus* | 3 | McAreavey, 1947 |
| Formicinae | *Pseudolasius* | *australis* | 0 | Emery, 1925 |
| Formicinae | *Pseudonotoncus* | *hirsutus* | 3 | Shattuck & O'Reilly, 2013 |
| outgroup | *Rhytidoponera* | *chalybaea* | 0 | AntWeb.org |
| Formicinae | *Rossomyrmex* | *anatolicus* | 3 | AntWeb.org |
| Formicinae | *Santschiella* | *kohli* | 3 | Forel, 1916 |
| Formicinae | *Teratomyrmex* | *greavesi* | 3 | Shattuck & O'Reilly, 2013 |
| outgroup | *Tetraponera* | *rufonigra* | 3 | Ward, 2001 |
| Formicinae | *Zatania* | *albimaculata* | 3 | AntWeb.org |

